# Structure and dynamics that specialize neurons for high-frequency coincidence detection in the barn owl nucleus laminaris

**DOI:** 10.1101/2022.07.22.501079

**Authors:** Ben Drucker, Joshua H. Goldwyn

## Abstract

A principal cue for sound source localization is the difference in arrival times of sounds at an animal’s two ears (interaural time difference, ITD). Neurons that process ITDs are specialized to compare the timing of inputs with submillisecond precision. In the barn owl, ITD processing begins in the nucleus laminaris (NL) region of the auditory brainstem. Remarkably, NL neurons are sensitive to ITDs in high-frequency sounds (kilohertz-range). This contrasts with ITD-based sound localization in analogous regions in mammals where ITD-sensitivity is typically restricted to lower-frequency sounds. Guided by previous experiments and modeling studies of tone-evoked responses of NL neurons, we propose NL neurons achieve high-frequency ITD sensitivity if they respond selectively to the small-amplitude, high-frequency fluctuations in their inputs, and remain relatively non-responsive to mean input level. We use a biophysically-based model to study the effects of soma-axon coupling on dynamics and function in NL neurons. First, we show that electrical separation of the soma from the axon region in the neuron enhances high-frequency ITD sensitivity. This soma-axon coupling configuration promotes linear subthreshold dynamics and rapid spike initiation, making the model more responsive to input fluctuations, rather than mean input level. Second, we provide new evidence for the essential role of phasic dynamics for high-frequency neural coincidence detection. Transforming our model to the phasic firing mode further tunes the model to respond selectively to the fluctuating inputs that carry ITD information. Similar structural and dynamical mechanisms specialize mammalian auditory brainstem neurons for ITD-sensitivity, thus our work identifies common principles of ITD-processing and neural coincidence detection across species and for sounds at widely-different frequencies.

**Author summary:** Differences in the arrival times of sounds at the two ears are essential for creating a sense of auditory space. For many animals, the utility of these interaural time-differences for sound source localization is thought to be restricted to relatively low-frequency sounds, due to limits of temporal precision in the auditory pathway. Barn owls, remarkably, use temporal processing to localize high-frequency (kilohertz-scale) sounds. This capability is critical for their activities as nocturnal predators. Building on insights from previous experimental and modeling studies, we propose that these neurons encode time differences in high-frequency sounds because they respond selectively to input fluctuations, and are relatively non-responsive to input mean. We use a biophysically-based computational model to show that electrical separation between a neuron’s input region (soma) and spike-generating region (axon) improves sensitivity to input fluctuations. This structural configuration produces linear integration of subthreshold inputs and rapid spike initiation, two dynamical features that improve time-difference sensitivity to high-frequency sound-evoked inputs. Neural coincidence detection in the neuron model is further enhanced if it operates in a phasic firing mode. Taken together, we provide new insights into the dynamical and structural mechanisms that support high-frequency sound localization by coincidence detector neurons.

## Introduction

A principal cue for sound source localization is the difference in arrival times of sounds at an animal’s two ears (interaural time difference, ITD). ITDs depend on animals’ head sizes and are small compared to typical neural time scales. The physiological ranges of ITDs are ±700 *µ*s in humans and ±170 *µ*s in barn owls, for instance. Neurons and neural circuits that process ITDs are specialized for temporal precision so that they can compare the timing of inputs at this submillisecond scale [1, 2]. Binaural ITD processing begins in mammals in the medial superior olive (MSO) and in birds in the nucleus laminaris (NL). Neurons in these two nuclei are often characterized as coincidence detectors because they respond with higher firing rates when brief inputs arrive nearly simultaneously [3–10] and because they have been considered as possible neural substrates for Jeffress’s influential theory of sound source localization by delay lines [11, 12].

Although MSO and NL neurons share a similar role in auditory processing, they operate at widely-different frequency ranges. ITD-sensitive MSO neurons are thought to primarily aid in the localization of low-frequency sounds due to the limits of phase-locking of their inputs [13] and consistent with the classical duplex theory of sound localization [14]. Phase-locking in the early auditory pathway of barn owls persists for much higher-frequency sounds [15]. Indeed, barn owls accurately localize sounds in the 4 kHz to 8 kHz range [16] and NL neurons shows ITD sensitivity in this frequency range as well [5, 17].

Extracting ITDs from kilohertz-scale signals poses a difficult computational challenge for coincidence detector neurons. At these high frequencies, inputs to NL neurons are not resolved as isolated synaptic events whose relative timing can be compared. Instead, sound-evoked votage-responses in the soma of NL neurons are small-amplitude oscillations at the frequency of the tone input, termed the sinusoidal analogue potential (SAP) [17]. Essential features of the SAP described by Funabiki and colleagues are 1) SAP amplitude varies with ITD, but the mean SAP level does not; and 2) NL firing rate increases with increases in SAP amplitude, but does not depend on SAP mean. These observations suggest that high-frequency ITD processing in the barn owl requires that NL neurons respond to fluctuations in their inputs (SAP amplitude) while remaining insensitive to slower changes in the baseline input level (SAP mean).

The presumption that NL neural firing should be selective for high-frequency fluctuations guides our analysis. Adapting a biophysically-based NL neuron model [17, 18], we identify structural and dynamical features that specialize NL neurons for high-frequency ITD sensitivity. We show that electrical separation of the soma and the axon improves neural coincidence detection. When the soma and axon regions are weakly-coupled, spike generation in the NL neuron model depends less on input mean than in the case of strong coupling between soma and axon. We identify two dynamical features associated with weak soma-axon coupling that benefit high-frequency neural coincidence detection: linear subthreshold dynamics (as opposed to amplified, supralinear responses to subthreshold inputs) and rapid spike initiation. The importance of electrical separation between soma and axon regions for high-frequency ITD processing accords with related studies of NL neurons [18, 19] (and similar work in MSO neurons [20, 21]), but we offer novel and clarifying insights into how weak soma-axon coupling produces fluctuation-sensitive dynamics that enhance ITD encoding.

Fluctuation-sensitivity is a feature of phasic neurons (also known as Type III excitability) [22–25]. Phasic neurons respond to a prolonged pulse of current (constant level) with a single spike at pulse onset, with no possibility for repetitive firing [26]. NL neurons exhibit phasic firing *in vitro* [7], but biophysically-based models developed previously for NL neurons are tonic (produce repetitive spiking in response to sufficiently strong constant inputs) [17, 18]. We show that transforming our model to the phasic firing further tunes the model to respond selectively to the small amplitude, high-frequency signals that carry ITD information. Thus we add to the understanding of the functional importance of phasic dynamics by showing phasicness beneftis high-frequency coincidence detection. In sum, our work points to coherent principles of soma-axon coupling configurations and spiking dynamics that specialize auditory neurons for coincidence detection across species and for sounds at widely-different frequency scales.

## Results

### Time-difference sensitivity is greater for weakly-coupled soma and axon regions

To study neural coincidence detection and ITD sensitivity in the barn owl NL, we constructed a two-compartment model with parameters adapted from an established model of these neurons [17]. The two compartments represent a soma-dendritic region (compartment 1) and an axonal region of spike generation (compartment 2). Synaptic inputs target the first compartment and spike-generating sodium currents are restricted to the second compartment with dynamics described by Hodgkin-Huxley-type nonlinear differential equations. We provide details in Methods. In NL neurons, spikes are thought to be primarily generated in the axon initial segment [18]. Axon structure and physiology can vary across the NL [19, 27]. Our first goal, therefore, was to systematically explore the effects of electrical coupling between soma and axon regions on firing rate responses to high-frequency inputs. We did this by parameterizing the NL model to describe the strength of electrical connection between the two compartments. We identified constants for forward coupling strength (*κ*_1*→*2_) and backward coupling strength (*κ*_2*→*1_) based on the method in [21]. These coupling constants can also be understood as the attenuation ratios of steady-state voltages.

By construction, all models (regardless of coupling configuration) have nearly identical passive soma dynamics, but the addition of sodium current in the second compartment leads to marked difference in spiking dynamics. In particular, models with weak soma-axon coupling have large-amplitude, fast-initiating, “all-or-nothing” spikes (Fig 1B) whereas spikes in strongly-coupled models are more graded with more gradual upstrokes (Fig 1D). To make consistent comparisons across coupling configurations, we separately determined maximal sodium conductance for each coupling configuration to achieve a fixed peak firing rate in simulated responses. Specifically, we generated two synaptic input streams (meant to represent 4 kHz tone-evoked responses from the two ears) with a possible time difference (ITD). For in-phase inputs (ITD= 0 ms), such as those shown in Fig 1A1, we set maximum sodium conductance so that the average firing rate of the NL neuron model was 500 spikes per second. Importantly, when synaptic events were evoked by out-of-phase 4 kHz sine waves (ITD= 125*µ*s, as in the example in Fig 1A2), NL firing rates decreased, indicatingITD-sensitivity. In these examples, the model with weaker coupling produces just two spikes (Fig 1B2) (a decrease from the in-phase response). The model with stronger coupling produces five spikes in response to the same input (Fig 1D2) which is nearly equal to its in-phase firing rate. The coupling configuration *κ*_1*→*2_ = 0.9 and *κ*_2*→*1_ = 0.5 is similar to the previously-developed model that is the starting point for our work [17]. We show responses of this model in Fig 1C.

**Fig 1.**
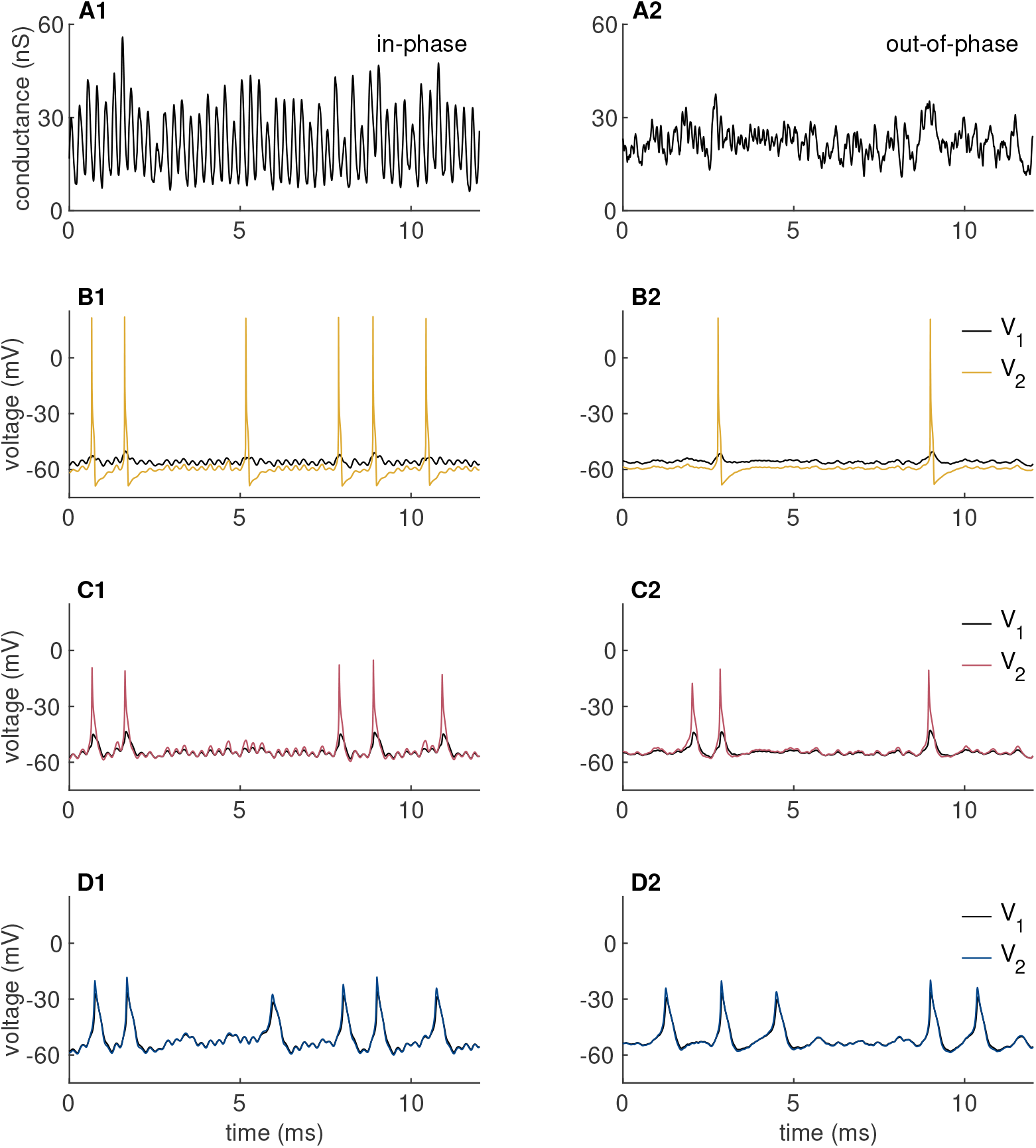
Spiking dynamics of a two-compartment NL model. (A) Synaptic conductance time-courses for 4 kHz in-phase (left column) and out-of-phase (right column) inputs. (B-D) Membrane voltage responses to inputs in (A), using three different soma-to-axon coupling configurations: (*κ*_1*→*2_, *κ*_2*→*1_) = (0.3, 0.2) in B, (0.9, 0.5) in C, and (0.9, 0.9) in D. Gray curves in these panels show soma voltage (*V*_1_) and colored curves show axon voltage (*V*_2_).

We next computed ITD tuning curves by simulating neural responses to inputs with a range of time differences (Fig 2A). For the strongly-coupled model, in which *V*_1_ and *V*_2_ are nearly isopotential, firing rates do not vary substantially with ITD. This is evident in the relatively flat firing rate curve for the case of *κ*_1*→*2_ = *κ*_2*→*1_ = 0.9 (Fig 2A). The other coupling configurations produce larger differences in the firing rates evoked by in-phase inputs versus out-of-phase inputs. This is our primary result – electrical separation of soma and axon improves ITD sensitivity in simulated NL responses.

**Fig 2.**
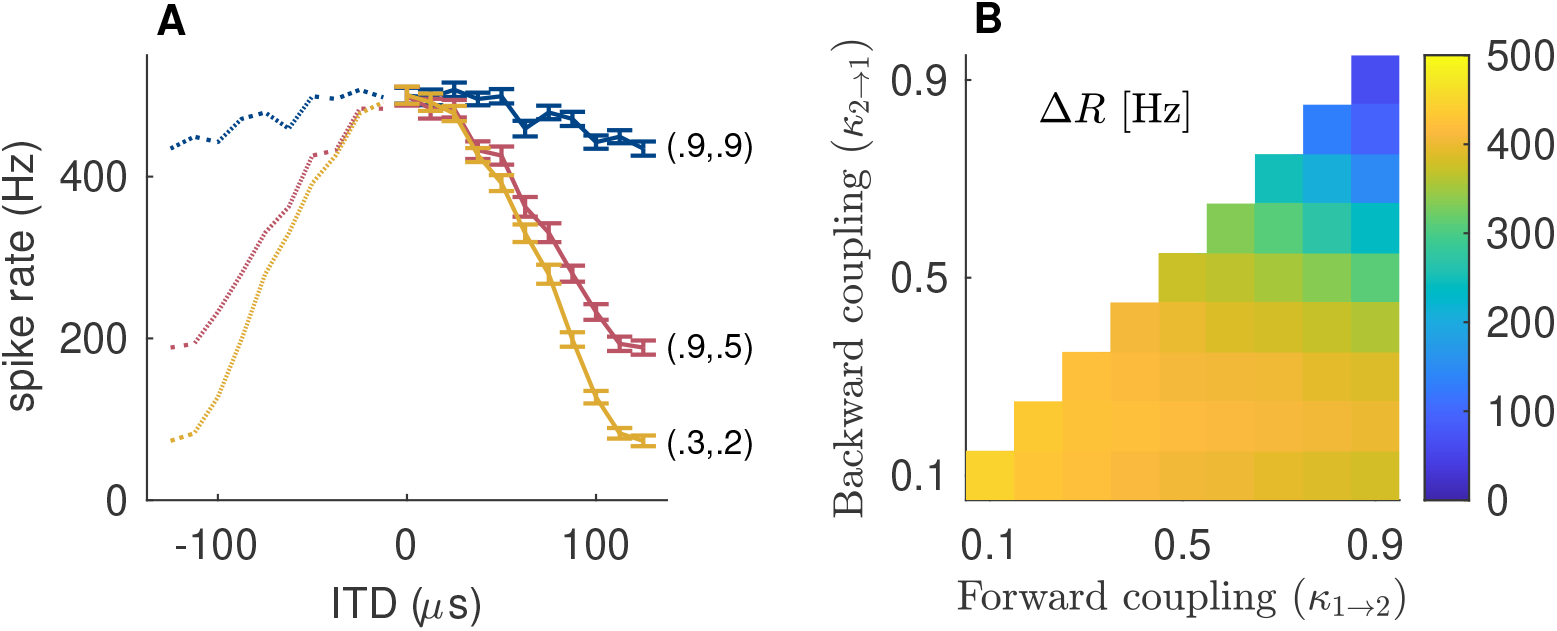
Tuning to input time difference requires electrically-isolated soma and axon compartments. (A) ITD tuning curves (spike rate as function of input time difference) in response to 4 kHz synaptic inputs. Coupling configurations and color scheme is same as Fig 1. Dotted lines are a reflected version of computed values. Error bars represent standard error in the mean firing rate from 100 repeated simulations. Sodium conductance is determined so 500 spikes/sec is peak firing rate for all configurations. (B) Difference between in-phase and out-of-phase firing rates across across the space of coupling configurations (Δ*R*, scale bar at right). Δ*R* is smallest for strong coupling and increases for electrically-isolated soma and axon compartments (weaker coupling).

To expand on these initial observations, we measured the difference between in-phase and out-of-phase firing rates across a full range of coupling configurations (Fig 2B). This measure of ITD tuning curve depth, which we denote by Δ*R*, is commonly-used in simulation studies and recordings of NL neurons [17, 18, 28]. By this measure, there is a relatively broad range of coupling configurations with large Δ*R* (good ITD sensitivity). The disadvantage of strong coupling is confined to a relatively small region of parameter space (small Δ*R* in the upper-right corner).

In the following, we explain this effect of coupling configuration on ITD sensitivity by clarifying the signal processing imperative for NL neurons to respond to input fluctuations, not mean level, and by explaining how structure creates dynamics that support effective high-frequency ITD coding.

### Coding imperative: ITD represented by input fluctuations

Informed by insightful experimental and theoretical work in the NL [17, 18, 29, 30], we take the view that ITD information is delivered to NL neurons via fluctuating synaptic inputs. These fluctuations evoke the small-amplitude, high-frequency oscillations in the somatic membrane potential of NL neuron termed the sinusoidal analogue potential (SAP) [17]. For NL neurons to effectively encode ITD signals, the amplitudes of input fluctuations should drive NL spiking activity, not the mean input level. An idealized view of this computation is NL neurons must monitor their synaptic inputs and use spike generation in the axon to signal when voltage fluctuations in the soma exceed some threshold. Threshold crossings should be determined by fluctuation amplitude and not mean input level.

We illustrate this coding perspective by computing, on a cycle-by-cycle basis, the mean and amplitude of the excitatory synaptic conductances that are inputs to the NL model. The period of each cycle is 250 *µ*s, as dictated by our use of a 4 kHz input tone. A scatter plot of 100 cycles of synaptic inputs reveals, as expected, that in-phase synaptic inputs are more distinguishable from out-of-phase inputs by differences in fluctuation amplitude, not the mean input level (Fig 3A, see also Fig 1A for example synaptic conductance time-courses).

**Fig 3.**
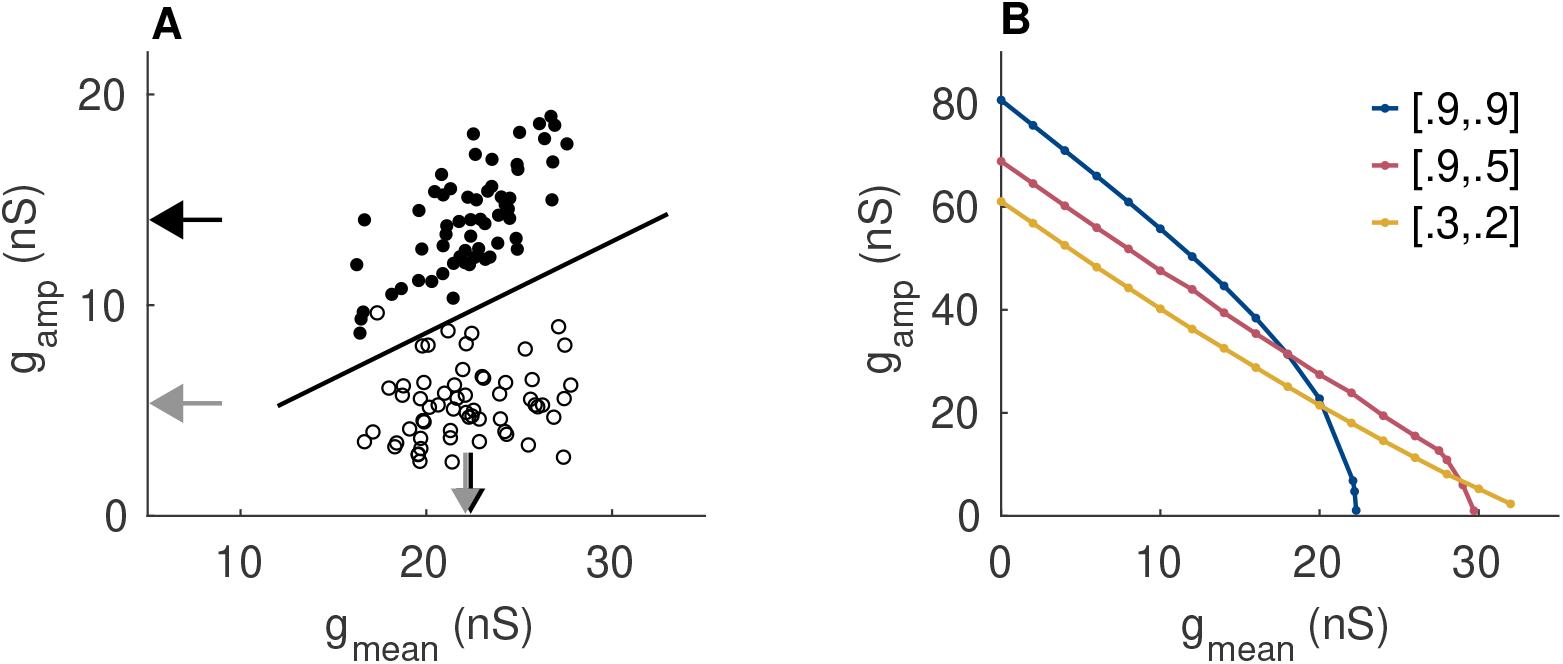
High-frequency ITD processing requires sensitivity to cycle-amplitude, not cycle-mean, of fluctuating inputs. (A) Scatter plot of mean and amplitude of synaptic conductance measured relative to period of 4 kHz input frequency for in-phase inputs (filled circles) and out-of-phase inputs (empty circles). Arrow heads indicate mean values of these measures (black arrow for in-phase, gray for out-of-phase). A classification boundary based on Fisher’s linear discriminant is indicated by the solid line. (B) Thresholds of repetitive firing in two-compartment NL model in response to sine-wave conductance. Coupling configurations and color scheme are same as Fig 1.

To explore further this idealized view of NL neurons as cycle-by-cycle observers of their inputs, we computed a classifier boundary defined using Fisher’s linear discriminant [31]. Synaptic inputs that fall above the boundary would be reported by an ideal observer of these synaptic conductances as in-phase inputs. The classifier boundary is upward sloping because fluctuation amplitudes are positively-correlated with mean input level for in-phase inputs. This suggests a stringent coding imperative for NL neurons: they should generate spikes when the amplitude of their high-frequency fluctuating inputs are sufficiently large and, also, they should avoid generating spikes to inputs with large mean values.

This description of an NL neuron as a signal classifier observing its inputs is, of course, an over-simplification of the biophysical processes at work. We do not expect NL neurons to act exactly like such ideal observers because intrinsic dynamics of NL neurons do not allow spike-generation to occur on a cycle-by-cycle basis in response to high-frequency inputs (due to refractory periods, for instance). Nevertheless, this perspective helps clarify which characteristics of NL excitability may enhance ITD sensitivity.

To relate this signal classification perspective to modeling results, we used sinusoidal conductance inputs (as opposed to the random synaptic inputs used in other simulations) to compute thresholds for repetitive firing as a function of input mean and amplitude (Fig 3B). For all coupling configurations, the slope of these threshold curves was negative, indicating the model becomes more excitable for larger mean input levels. This is inconsistent with ideal observer’s classification boundary. Among these coupling configurations, though, the strongly-coupled model had the steepest threshold curve. For *g*_*mean*_ larger than roughly 15 nS, changes in mean input level alone, not fluctuation amplitude, can drive changes in firing rate for this coupling configuration. This parallels the simulated ITD tuning curves presented above (Fig 2D) by showing that strong electrical coupling between soma and axon is disadvantageous for ITD tuning because configurations with electrically-separated soma and axon compartments are more selective for input fluctuations (rather than input mean).

### Distinctive dynamics of strongly-coupled models that hinder high-frequency coincidence detection

What accounts for enhanced ITD-tuning for models with stronger soma-axon coupling? We identified two dynamical features caused by strong coupling that distinguish those models from models with electrical isolation between soma and axon. These features are: 1) supralinear subthreshold integration in the soma and 2) slow spike initiation in the axon. By supralinear integration, we mean that the subthreshold current-voltage relation is nonlinear with positive concavity as in the *I*-*V* curves shown in Fig 4A. Amplification of subthreshold voltage is greatest for models with strong soma-axon coupling because strong soma-to-axon (forward) coupling activates sodium conductance and, in turn, strong axon-to-soma (backward) coupling enables axonal sodium current to depolarize the soma.

**Fig 4.**
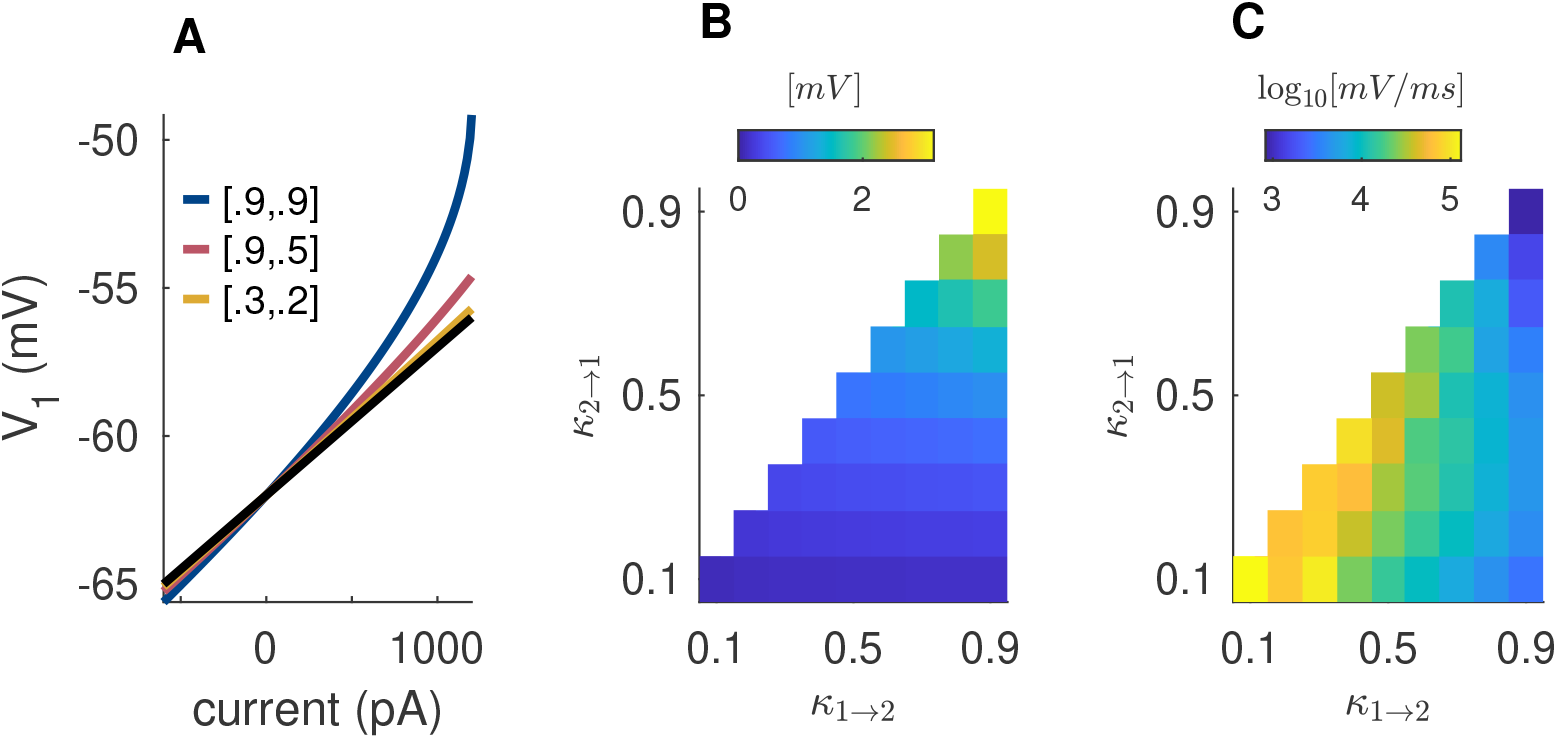
Nonlinear mechanisms that distinguish strongly-coupled models from models with electrically-isolated compartments. (A) Stready-state soma voltage response to constant input current (*I*-*V* relation). Color code and coupling configurations are same as Fig 1. Black line is response of the model *g*_*Na*_ removed (linear response with 5 *M*Ω soma input resistance). (B) Subthreshold *V*_1_-amplification measured as the difference between steady-state *V*_1_ responses to 1000 pA inputs for models with and without sodium current. (C) Maximum *V*_2_ slope during spike upstrokes, calculated as average of 100 responses to in-phase inputs. Colors given on a logarithmic scale as indicated.

We quantified the amount of amplification across the range of coupling configurations by calculating the steady state *V*_1_ (soma) voltage response to a subthreshold input (1000 pA constant current) and compared it to the *V*_1_ value that would be expected for a purely passive model (recall that passive soma input resistance is fixed as a constant for all coupling configurations). The largest amplification occurs in models with strong soma-axon coupling and models with weak soma-axon coupling have nearly linear *I*-*V* relations (Fig 4B).

In addition to affecting subthreshold integration, coupling configuration also changes the shape and dynamics of spikes (recall Fig 2A,B). In strongly-coupled models, voltage dynamics in the two compartments track one another closely. Spikes occur in both compartments with *V*_1_ dynamics slowing the rate of upstroke. In weakly-coupled models, by contrast, voltage dynamics in the axon are insulated from the soma. In these cases, fast activation of sodium current during a spike increases *V*_2_ rapidly and without hindrance from the more slowly-depolarizing soma.

We measured the maximum slope of spike upstroke in *V*_2_ across coupling configurations (Fig 4C). The rates of spike upstrokes in weakly-coupled models are as much as one hundred times faster than those in the most strongly-coupled model. This finding is consistent with previous modeling work showing that moving the location of a localized spike initiation zone to positions on the axon more distant from soma leads to steeper spike upstrokes [32].

### Advantages of linear subthreshold integration and fast spike initiation illustrated in an integrate-and-fire model

We hypothesized that the two dynamical features identified above that distinguish strongly-coupled models from other coupling configuration (supralinear subthreshold integration and slow spike initiation) could account for degraded ITD sensitivity in models with strong soma-axon coupling. Our reasoning was as follows. Inputs to NL neurons are relatively weak — the SAP (soma voltage fluctuations) is on the order of a few millivolts [17]. Models of NL neurons must operate, therefore, near threshold so these small-amplitude fluctuations suffice to evoke spiking in NL neurons [17, 18]. At the same time, to maintain high-frequency ITD sensitivity, NL neurons must not respond to changes in input mean. Nonlinear amplification of subthreshold inputs are detrimental for high-frequency coincidence detection, therefore, because constant or slowly-varying subthreshold inputs that are nonlinearly amplified can drive spiking activity even in the absence fluctuations related to meaningful ITD information. There is greater amplification of soma voltage in strongly-coupled models than models with weak axon-to-soma coupling, so the strongly-coupled configuration is not optimal for preventing spike initiation in response to constant or slowly-varying inputs.

The second distinctive feature of strongly-coupled models is (relatively) slow spike initiation. Fast spike initiation should be beneficial for high-frequency ITD detection. The larger-amplitude portions of high-frequency fluctuating inputs represent coincident inputs corresponding to salient ITD information. When a brief but strong input fluctuation arrives (viewed as a small portion of a high-frequency oscillating input), a rapidly-initiated spike can be triggered before the input level decreases (during the next phase of the input oscillation). Neurons with slowly-initiating spikes, by contrast, may need to “integrate” over larger portions of their inputs before spikes fully develop. Neurons with gradual spike upstrokes would be less responsive to the brief and strong input fluctuations created by coincident inputs.

We illustrate how these two features affect high-frequency fluctuation sensitivity using an integrate-and-fire model customized with parametric control over subthreshold dynamics and spike upstroke speed. An interpolation parameter *p* switches the model between linear subthreshold dynamics (*p* = 0) and supralinear subthreshold dynamics (*p* = 1). When the dynamical variable in the model exceeds a threshold level, the dynamics change to exponential growth. A gain parameter *q* controls the exponential rate-of-rise of *x*. When *x*(*t*) exceeds the maximum value of *x*_*max*_ = 50 the dynamical variable is returned to *x*(*t*) = -5, a reset condition typical for representing spikes in integrate-and-fire model. See Methods for details and Fig 5A shows the graph of the piecewise-function that governs the intrinsic dynamics of the model. A linear subthreshold model with slow spike initiation is in Fig A1 (*p* = 0, *q* = 1) and a model with suprathreshold amplification and fast spike initiation is in Fig A2 (*p* = 1, *q* = 4). We simulated responses of this model to high-frequency sinusoidal inputs, varying input mean (*g*_*mean*_) and input amplitude (*g*_*amp*_). Examples of spiking responses are shown in Fig 5B with steeper spike upstroke visible in Fig 5B2 due to the larger value of *q* in that simulations.

**Fig 5.**
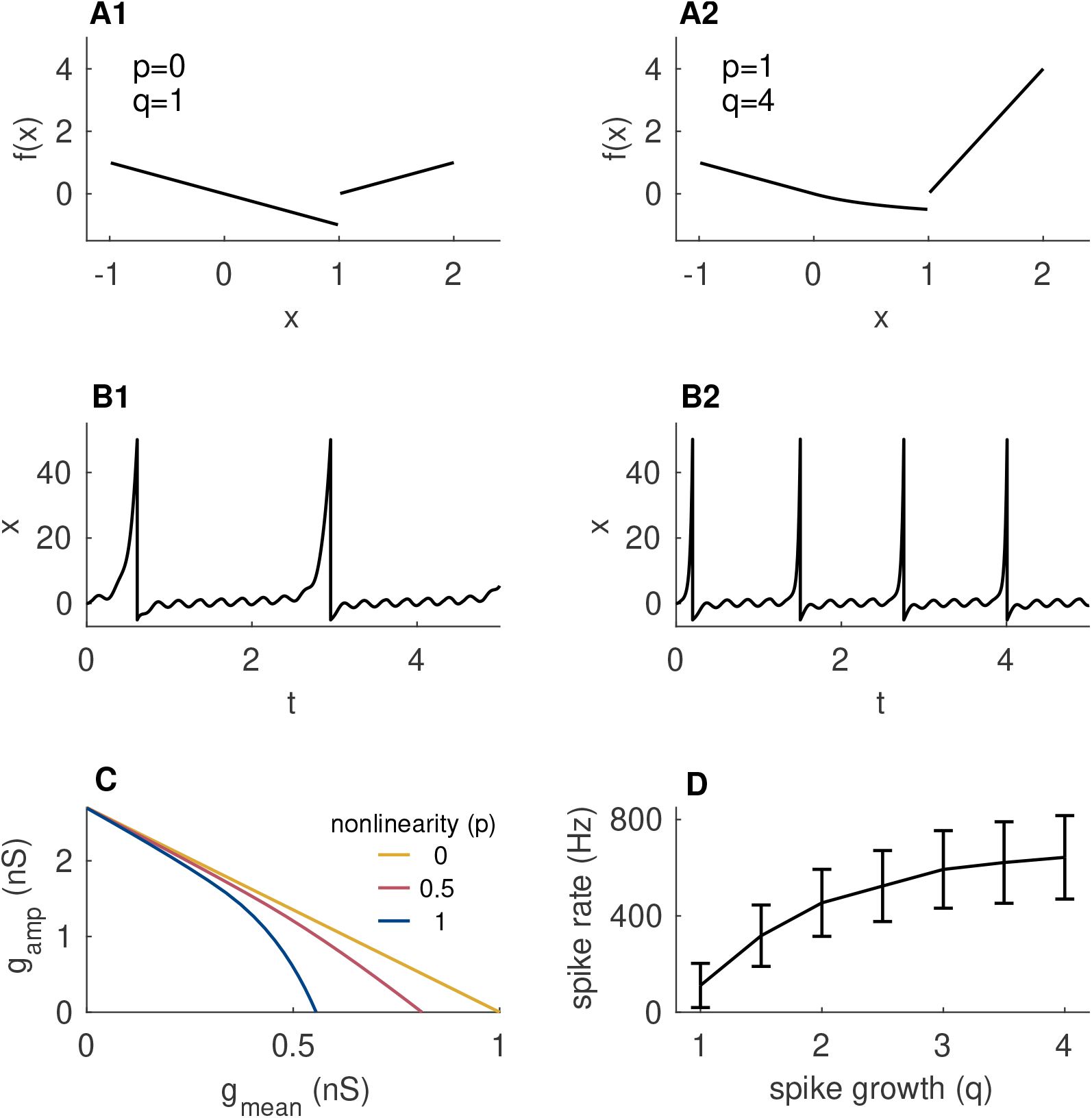
Nonlinear integrate-and-fire model illustrates effects of subthreshold amplification and spike initation speed on sensitivity to high-frequency fluctuations. (A) Piecewise-defined function that governs model dynamics. An interpolation parameter (*p*) controls whether subthreshold integration is linear (*p* = 0 in A1) or supralinear (*p* = 1 in A2). A second parameter (*q*) controls the speed of spike initiation (the slope of *f* (*x*) where *x >* 1). (B) Time-courses of the dynamical variable *x*(*t*) in response to 4 kHz sine-wave input, with parameter sets in B1 and B2 corresponding to those in A1 and A2. (C) Thresholds for repetitive firing in response to 4 kHz input with varying mean (ordinate) and amplitude (abscissa). The subthreshold nonlinearity *p* affects the slope of these threshold curves (as shown), but the spike slope *q* does not. (D) Mean spike rate in response to 4 kHz inputs with fluctuating cycle-amplitudes (with zero mean input) and linear subthreshold dynamics (*p* = 1). Error bars show standard deviations of 500 repeated trials.

We have parametric control over subthreshold amplification and rate of spike initiation in this integrate-and-fire model. We first computed thresholds for repetitive spiking as a function of input mean and amplitude and found that increasing the degree of subthreshold amplification (increasing *p*) produces threshold curves that slope downward more steeply (Fig 5C). This is qualitatively consistent with repetitive spiking threshold curve for the strongly-coupled two-compartment model (Fig 3B). In the simulations shown, we used *q* = 3 for the rate of spike growth in the integrate-and-fire model. In additional simulations (not shown) we varied *q* between 1 and 5 and observed no changes to these threshold curves.

Next, we varied the spike growth parameter *q* to confirm that rapid spike initiation increased sensitivity to high-frequency fluctuations (Fig 5D). For these simulations we used inputs that were sinusoidal, but with oscillation amplitude that varied randomly on a cycle-by-cycle basis. Our rationale for this input structure was that most cycles would be subthreshold oscillations (by design) with some larger-amplitude fluctuations occurring at random. Higher firing rates in response to this random input would indicate greater sensitivity to the brief but large fluctuations produced by coincident inputs. We found, as expected, firing rates increased with the speed of spike initiation. The results shown in Fig 5D are for *p* = 0 (linear model), and we found qualitatively similar outcomes for models with nonlinear subthreshold dynamics (*p >* 0, results not shown).

### Phasic dynamics enhance high-frequency ITD-sensitivity

To this point we have illustrated now soma-axon electrical separation promotes fluctuation-sensitive dynamics that are beneficial for high-frequency ITD sensitivity. A well-known mechanism for fluctuation-sensitivity in neurons is phasic dynamics. NL neurons display phasic dynamics *in vitro* [7] and phasic neurons are known to respond selectively to input fluctuations (and not input mean) [22, 23, 26]. Previous models of NL neurons have not exhibited phasic firing [17, 18], so we were compelled to explore whether phasic dynamics would enhance high-frequency ITD sensitivity in the NL neuron model. We began by inspecting the dynamics of a reduced version of the model in which soma voltage *V*_1_ is a (constant) input strength, sodium activation is set instantaneously to its voltage-dependent steady state value *m*_*∞*_(*V*_2_), and *g*_*KHT*_ set to 0. These manipulations yielded a two-variable model (*V*_2_-*h*) of axonal dynamics. The phase plane for this reduced axon model contains a fixed point that loses stability as it transitions from the left branch of the *V*_2_-nullcline to the middle branch for sufficiently large input strength *V*_1_ (Fig 6A1). This transition is characteristic of tonic dynamics [33].

**Fig 6.**
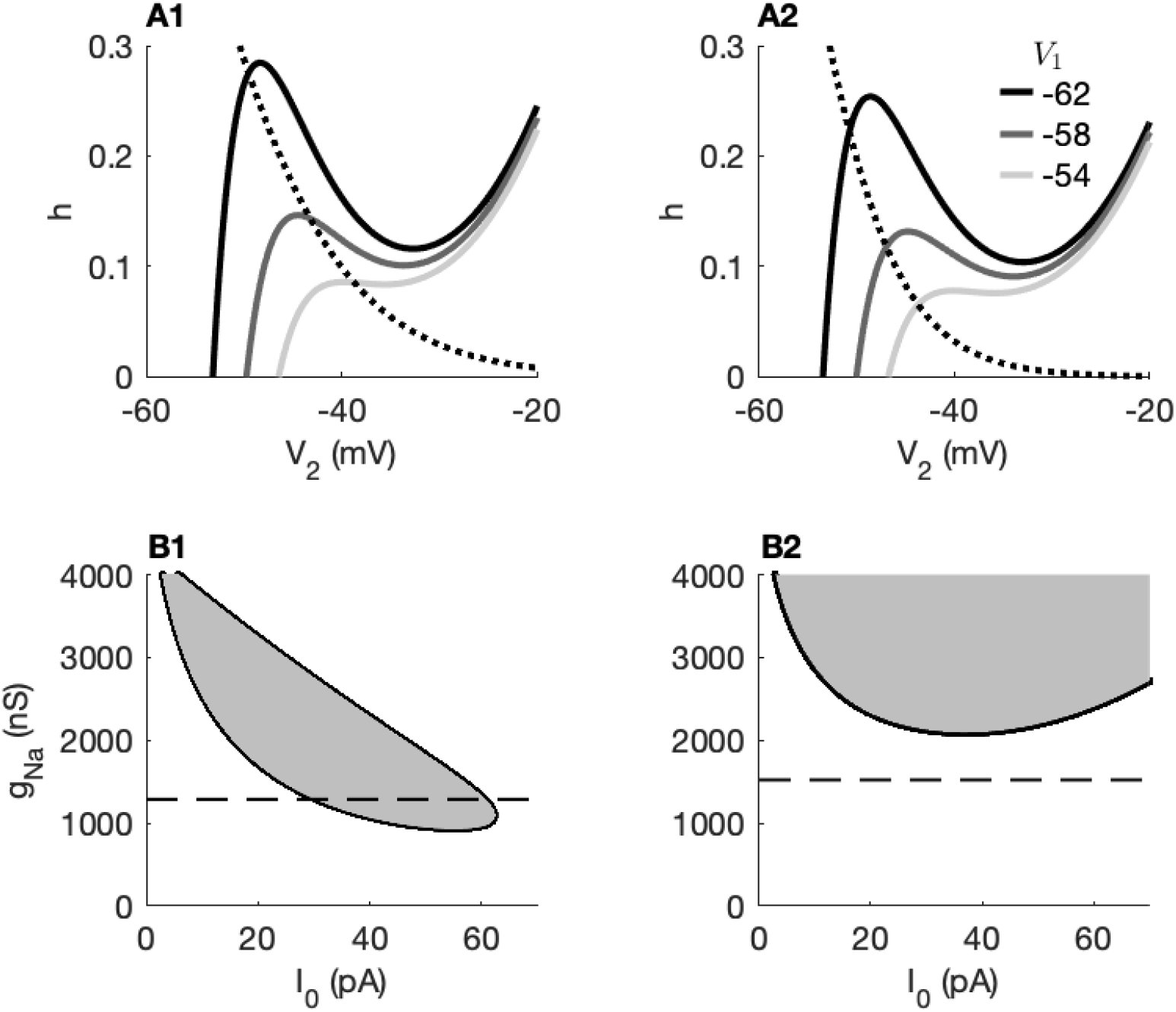
Modification of sodium inactivation converts two-compartment model from tonic to phasic firing. Coupling configuration is *κ*_1*→*2_ = 0.9, *κ*_2*→*1_ = 0.5. (A) Phase-plane diagrams for two-variable (reduced) axon model. Dotted line shows *h*-nullcline for the control model (*σ* = 7.7 in A1) and a model with steeper *h*_*∞*_ function (*σ* = 5 in A2). Solid lines show *V*_2_-nullclines for varying *V*_1_ (treated as a constant, input parameter). (B) Two-parameter bifurcation study of two-compartment (full) NL model showing combinations of constant input current (*I*_0_) and sodium conductance (*g*_*Na*_) that produce tonic firing (shaded region) or phasic firing (non-shaded region). Horizontal dashed line marks the *g*_*Na*_ value used in ITD simulations (to satisfy criterion of 500 sp/s peak firing rate). Values of *σ* (steepness of *h*_*∞*_ in B1 and B2 correspond to values in A1 and A2, respectively.

From this geometric approach, we observed that a convenient way to convert this two-parameter axon model to the phasic firing mode was to steepen the sodium inactivation function *h*_*∞*_ by decreasing the parameter *σ* in Eq 3. The default value used in Fig 6A1 and based on previous models [17] is *σ* = 7.7. When we steepened sodium inactivation by setting *σ* = 5, we found the fixed point remained stable and located on the left-branch of the *V*_2_-nullcline (Fig 6A2). This indicates no possibility of repetitive firing to constant inputs [26].

We next confirmed that this manipulation of *h*_*∞*_ acted similarly in the full two-compartment NL model. For various values of sodium inactivation steepness (*σ*), we performed a two-parameter bifurcation analysis using input current strength *I*_0_ and sodium conductance *g*_*Na*_ as bifurcation parameters (Fig 6B). For the reference model using *σ* = 7.7 a region of tonic firing was present for sufficiently strong input current (gray shaded region in Fig 6B1). Maximal sodium conductance for this configuration was *g*_*Na*_ = 1286 nS (marked by dashed line that passes through the tonic firing region). This model fires repetitively for inputs with sufficiently large mean values. The model with steeper *h*_*∞*_ and *g*_*Na*_ = 1522 nS does not fire repetitively to constant inputs, regardless of the input level (Fig 6B2). Recall that *g*_*Na*_ is different in these two models because we selected *g*_*Na*_ separately for all model configurations to maintain the 500 sp/s firing rate at 0 *µ*s ITD.

Converting the model from tonic to phasic firing substantially altered sensitivity to synaptic fluctuations and improved ITD tuning. We first computed thresholds for repetitive firing to sinusoidal conductance (Fig 7A). Earlier we observed these curves were downward sloping (Fig 3B), even though an idealized signal classification view of this problem indicated that upward-sloped threshold curves would be optimal for coincidence detection (Fig 3A). Reducing *σ* had the effect of flattening these threshold curves so that the model could be more sensitive to input fluctuations and less sensitive to input mean. In fact, for the steepest *h*_*∞*_(*V*) curve tested (*σ* = 3, also a phasic model), portions of the threshold were upward-sloped and thus more similar to the ideal observer’s classification boundary.

**Fig 7.**
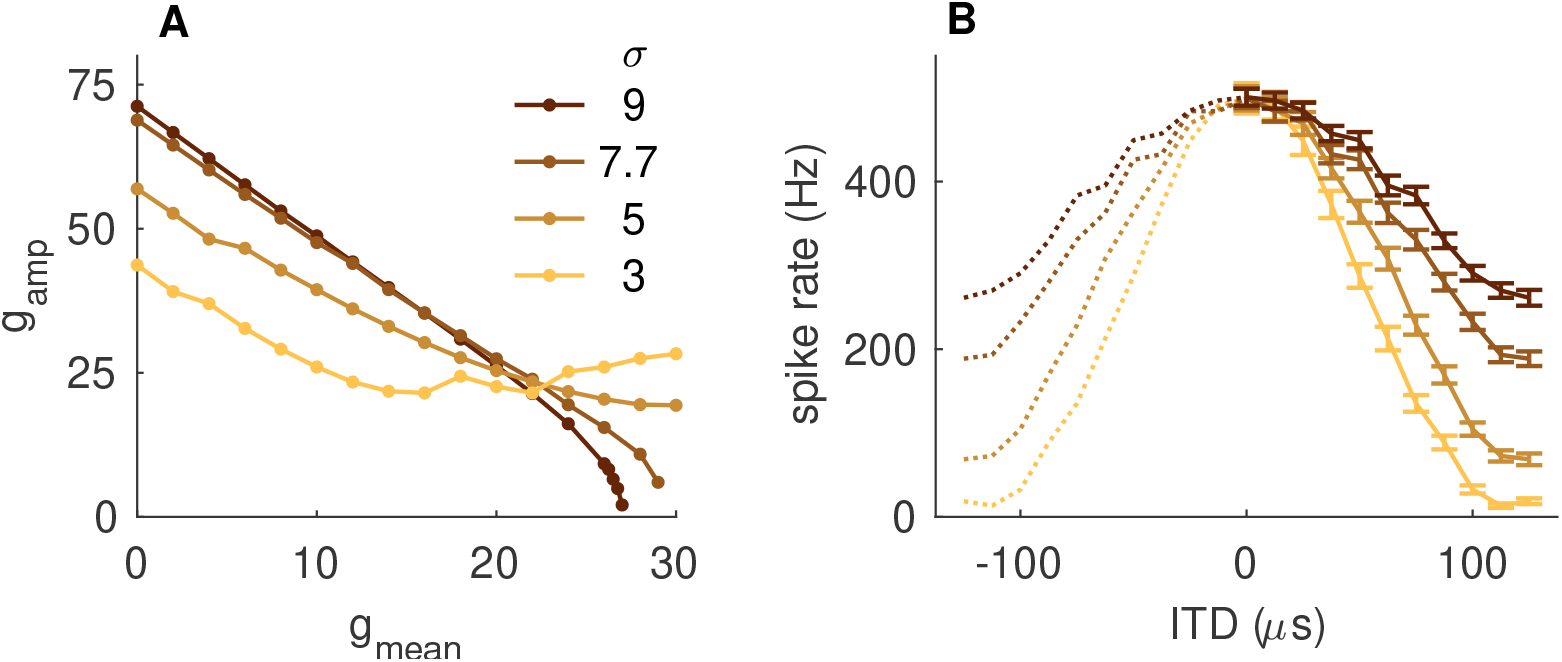
Phasic dynamics enhance fluctuation sensitivity and ITD tuning in the two-compartment NL model. (A) Thresholds for repetitive firing in response to 4 kHz sinusoidal conductance with varying mean (ordinate) and amplitude (abscissa). Compare to Fig 3B and observe the slope of the threshold curves is less step, and can even turn positive, for phasic firing models (smaller *σ*). (B) ITD tuning curves (spike rate as function of input time difference) in response to 4 kHz synaptic inputs. Format is same as Fig 2A. Tuning curves narrow for smaller *σ* demonstrating advantage of phasic firing for high-frequency ITD processing.

Consistent with these changes in responsiveness to sinusoidal conductance, we found that converting the model to phasic-firing enhanced ITD tuning (Fig 7B). Visible changes to ITD tuning curves for smaller *σ* include both narrower tuning curves and greater peak-to-trough differences (Δ*R*). For the default parameter value of *σ* = 7.7 (tonic mode) the model fired vigorously to out-of-phase inputs (200 spikes/second, approximately). Out-of-phase firing is nearly extinguished for sufficiently steep sodium inactivation (*σ* = 3, phasic firing mode) while maintaining the criterion level of excitability to in-phase inputs (500 spikes/second).

## Discussion

Temporal precision is a hallmark of neural processing in the auditory pathway [13, 15, 34–37]. Sound localization on the basis of interaural time differences (ITDs) is one of the most temporally-demanding aspects of auditory perception. Sound source location is encoded (in part) by submillisecond-scale time differences in sounds arriving at the two ears. Unraveling the physiological specializations of temporally-precise binaural neurons has been the subject of sustained investigation. Features that specialize these neurons to act as temporally-precise coincidence detectors include dendritic integration [28, 34, 38–40], ionic currents active at subthreshold voltages [9, 25, 41], synaptic inputs [42–46], and spike generator regions in the axon at locations remote from the soma [18–21, 27].

### Electrical separation of the soma and axon is essential high-frequency coincidence detection

We focused our attention on the dynamics of spike generation and the nature of soma-axon coupling since fine-tuning of soma-axon coupling may be particularly relevant for high-frequency ITD processing. In the nucleus laminaris (NL), where neurons can face the extreme challenge of extracting ITD information from kilohertz-scale inputs, previous work has shown that spike initiation zones in the axon are more distant and smaller in size for neurons that have higher characteristic frequencies [19]. Computational modeling also showed that ITD sensitivity is improved if spike initiation occurs only in the axon, with the soma structured as large and passive (without sodium current) [18]. Similar structural advantages may help MSO neurons operate at the upper frequency-limit of their ITD sensitivity [20, 21, 47].

A typical explanation for the advantage of remote spike initiation zones in NL neurons is that this configuration can prevent temporally-summated inputs in the soma from causing sustained depolarization of the axon. Insulating the spike-generator region from sustained depolarization prevents inactivation of sodium channels that would suppress excitability [19, 27]. In addition, impedance analysis in a NL model indicates that the transfer of high-frequency voltage oscillations from soma to axon is greater if sodium current is absent from the soma region [18]. Synthesizing these arguments, one could conclude that the NL soma-axon connection should act as a high-pass filter. There should be minimal attenuation (or even amplification) in the transfer of high-frequency voltage fluctuations to the axon, but mean soma voltage or slow variations in soma voltage should not pass to the axon.

Using a signal detection analogy, we considered the same ITD encoding challenge, but from the point of view of using axonal spiking as a means to monitor NL inputs for ITD information. Informed by current understanding of NL responses to high-frequency pure tone inputs (the sinusoidal analogue potential, SAP, as measured in [17] and studied further in [29, 30]), our view is that spike generation in NL neurons should be insensitive to slowly-varying changes in inputs (the kinds of changes that would be associated with temporal summation of non-coincident inputs). Instead, NL neurons should fire in response to the high-frequency fluctuations that are evoked by coincident synaptic events. Said differently, we concur that NL neurons should have high-pass-like behavior to be effective ITD processors, but we locate mechanisms for high-pass-like behavior in several nonlinear aspects of NL neural dynamics.

We showed that high-frequency ITD sensitivity is severely degraded by strong electrical coupling between the soma and spike-generating regions (Fig 2). Strong soma-axon coupling allows for sodium in the axon to act as a source of nonlinear amplification of subthreshold inputs in the soma (Fig 4A,B). This causes neural excitability to depend on mean input level (Fig 3B). Structural configurations with weaker soma-axon coupling linearly integrate subthreshold inputs and thus respond more selectively to input fluctuations. This is consistent with a previous finding that a passive soma enhances coincidence detection in an NL model because sodium in the soma nonlinearly-amplifies subthreshold inputs [18]. In addition, strong coupling slows the speed of spike initiation (which we measured as the slope spike upstroke, Fig 4C). Similar observations about the relationship between spike upstroke and soma-axon coupling have been made previously [32]. Fast spike initiation aids in responding to high-frequency input [47–49], so this is a second reason why electrical separation of the soma from the axon enhances high-frequency ITD sensitivity.

If insulation from temporally-summating inputs (to prevent sodium inactivation in the axon) were the primary benefit of weak soma-to-axon coupling, one might expect that ITD sensitivity should decrease in our model with increases in the forward-coupling constant. Some minor trends to this effect are evident in Fig 2B for configurations with strong backward coupling. For weaker backward coupling, though, ITD sensitivity does not depend on forward coupling strength. Recall, though, that we selected sodium conductance separately for each coupling configuration to maintain a comparable degree of excitability (500 sp/sec to in-phase inputs). For instance, maximum sodium conductance is *g*_*Na*_ = 4304 nS for the (*κ*_1*→*2_, *κ*_2*→*1_) = (0.3, 0.2) configuration and is *g*_*Na*_ = 428 nS for the (*κ*_1*→*2_, *κ*_2*→*1_) = (0.9, 0.2) configuration. Reduced sodium conductance for increases in forward coupling strength may explain why we did not find that strong forward coupling, on its own, degrades ITD sensitivity by inactivating sodium currents.

### Phasic excitability, a generic mechanism for neural coincidence detection, also improves high-frequency ITD processing

NL neurons exhibit phasic firing *in vitro* [7], but the Hodgkin-Huxley-type models previously developed for NL neurons do not [17, 18]. We determined that our NL model became more sensitive to input fluctuations and less sensitive to input mean when it was converted to the phasic firing mode (Fig 7A). In addition, phasic firing resulted in improved ITD tuning (Fig 7B).

We steepened the sodium inactivation steady-state curve *h*_*∞*_(*V*) as a straightforward way to convert the model to phasic firing (see also [26]). A previous study of coincidence detection and temporal precision in MSO neurons left-shifted the *h*_*∞*_(*V*) curve as a means to toggle between phasic and tonic firing [25]. Either manipulation of *h*_*∞*_(*V*) creates phasic firing dynamics because they strengthen the negative-feedback effect produced by sodium inactivation at subthreshold voltage levels [25].

If a model is in a tonic firing mode then it can be tipped into a repetitive firing pattern by temporal summation of high-frequency inputs. This can be problematic for modeling NL neural activity because voltage-responses in the soma that trigger spikes are small-amplitude events (SAP fluctuations on the order of a few millivolts) [17]. Sodium current in these models must be carefully calibrated, therefore, so that small fluctuations can evoke spikes but temporal summation of inputs do not [28]. In past modeling studies, mean input levels have been selected to position the dynamics near the boundary of critical points for repetitive firing [18] or have incorporated synaptic suppression to reduce temporal-summation of synaptic inputs [17]. Some amount of fine-tuning is necessary for these neurons because of the nature of their synaptic inputs and the temporally-demanding computation they perform. Indeed, developmental and homeostatic processes do regulate axon physiology in NL neurons and their inputs [19, 27]. That being said, phasic excitability offers a robust mode for neurons to selectively respond to input fluctuations and remain insensitive to temporally-summated, slowly-varying input currents [22, 23, 26].

### Toward a unified view of binaural coincidence detector neurons

We have found ITD sensitivity to be enhanced if soma and axon regions are electrically separated in an NL neuron model and that phasic dynamics further enhance the function of these neurons as temporally-precise coincidence detectors. Similar structural configurations may enhance ITD processing by MSO neurons in mammals [20, 21, 47], suggesting common dynamic principles at work in these different neural systems. While MSO and NL neurons operate at different frequency scales and in the context of cross-species differences in their auditory pathways, they appear to share many physiological features. We have remarked that both MSO and NL neurons exhibit phasic firing. In addition, they are both characterized by several related physiological features (low input resistance, fast membrane time constant, prominent voltage-gated currents active at subthreshold voltages). There are, however, notable differences in these two circuits. See [50, for review], including discussion of the differences in the numbers and types of synaptic inputs. Continued explorations of the similarities and differences between these centers for binaural coincidence detection may clarify the function of MSO neurons with high characteristic frequency [51, 52] and may also provide insights into how ITD information can be delivered with the high-frequency stimuli used in cochlear implant technology [53, 54].

There are some features of binaural neurons and circuits that we have not included in our model but that can be understood in relation to our findings. Inhibitory feedback from the superior olivary nucleus improved ITD sensitivity in a model of NL circuit in chicken [55]. Inhibitory feedback that stabilizes mean input level (counteracting sustained depolarization due to temporal summation) would help the NL circuit transmit ITD information via input fluctuations. The function of inhibition in MSO neurons continues to be studied [44, 56–58] with some proposals that precisely-time inhibitory inputs shift the peaks of ITD tuning curves [42, 43, 59, 60] (but see also [10, 46]). Low-threshold potassium current is prominent in NL neurons and enhances ITD sensitivity in MSO neurons. As discussed by Ashida and colleagues, low-threshold potassium current renders their model “more tolerant to changes in DC amplitude” [18]. In other words, the negative-feedback effect of this current (active at subthreshold voltage levels) can prevent supralinear amplification of subthreshold inputs (and recall we found such amplification be detrimental to high-frequency ITD sensitivity).

In sum, the remarkable temporal precision of binaural coincidence detector neurons requires numerous cellular and circuit-based specializations. Tracing cross-species similarities between MSO and NL neurons provides useful perspectives on both systems. We have emphasized that the nature of high-frequency synaptic inputs requires NL neurons to respond selectively to fluctuations amplitude, not mean input level. Low-frequency ITD encoding in the MSO, in contrast, requires slope-sensitive neurons to respond selectively to a few well-timed inputs [9, 26, 42, 51]. Taken together, our findings add to the evidence that there are shared structural and dynamical principles underlying the encoding of sound source location by neural coincidence across different species and widely-different frequency ranges.

## Methods

### Two-compartment NL neuron model

We studied spiking dynamics of a barn owl NL neuron using a two-compartment model that had been developed previously [17, 18]. The model consists of a compartment with passive dynamics (compartment 1 with voltage variable *V*_1_, representing the soma region) and a compartment with excitable dynamics (compartment 2 with voltage variable *V*_2_, representing a spike-initiating node in the axon). Voltages in the two compartments are governed by coupled differential equations

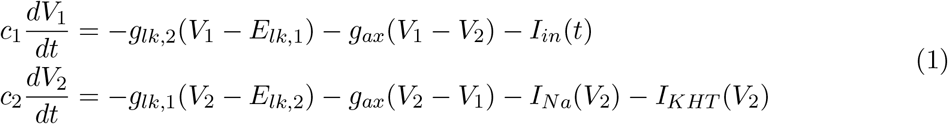

where capacitance (*c*), leak conductance (*g*_*lk*_), and leak reversal potential (*E*_*lk*_) can take different values in each compartment. The coupling conductance *g*_*ax*_ is the Ohmic coupling between the two compartments. The input current to the soma *I*_*in*_ typically represents either sinusoidal input current or conductance-based synaptic inputs. Voltage-gated ionic currents in the axon are spike-generating sodium current (*I*_*Na*_) and high-threshold potassium current *I*_*KHT*_. More details regarding these currents are given below.

### Passive parameters determined by soma-axon coupling

Following the method described in [21], we set passive parameter values so that *V*_1_ dynamics reproduced basic, physiologically-measurable properties of NL neurons (resting potential, input resistance, and membrane time constant). We then created a two-parameter space described by the strength of forward and backward couplings between the two compartments. With this approach we could study ITD sensitivity while systematically varying soma-axon coupling configurations [21, 61].

We set passive parameter values using values of three physiological constants similar to what has been reported in a previous studies of NL neurons [17]. These are input resistance in the soma (*R*_1_ = 5 *M* Ω), resting potential in the soma (*E*_*rest*_ = −62 *mV*, also used for *V*_2_ resting voltage), and soma membrane time constant describing the time-scale of exponential decay of *V*_1_ (*τ*_*exp*_ = 0.1 ms). We assumed the surface area of the first compartment is orders of magnitude larger than the surface area of the second compartment (2400 *µm*^2^ compared to 20 *µm*^2^), consistent with previous NL modeling studies following [17, 18].

We followed the approach in [21] to determine passive parameters in the two-compartment model. This enables systematic variation of coupling configuration will maintaining nearly identical passive *V*_1_ dynamics. Defining *U*_1_ and *U*_2_ to be the deviations of voltages from rest and removing the voltage-gated currents *I*_*Na*_ and *I*_*KHT*_, the passive dynamics relative to rest are:

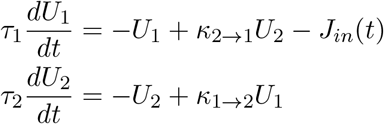

where the time constant parameters *τ*_*i*_ = *c*_*i*_*/*(*g*_*i*_ + *g*_*ax*_) for *i* = 1, 2 and *J*_*in*_ = *I*_*in*_*/*(*g*_1_ + *g*_*ax*_) is a rescaled input term. The soma-to-axon (forward) coupling parameter *κ*_1*→*2_ and axon-to-soma (backward) coupling parameter *κ*_2*→*1_ describe the impact of voltage deviations in one compartment on the other. Alternatively, these parameters can be thought of as steady-state attenuation factors between the two compartments. They are

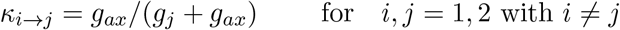

Due to the large discrepancy in membrane surface areas, there is a time-scale separation between the passive dynamics in the two-compartments (*V*_2_ is fast relative to *V*_1_). In particular, the ratio of time constants is *τ*_2_*/τ*_1_ = *ακ*_1*→*2_*/κ*_2*→*1_. For *α* = 20*/*2400 (as specified above) and the range of coupling constants used in our study, we have that *τ*_2_ takes values approximately 10 to 100 times smaller than *τ*_1_. Due to this separation of time-scales, we could use fast-slow analysis to uniquely define combinations of passive parameters that vary soma-axon coupling while maintaining nearly identical passive dynamics in the soma compartment (see [21] for details):

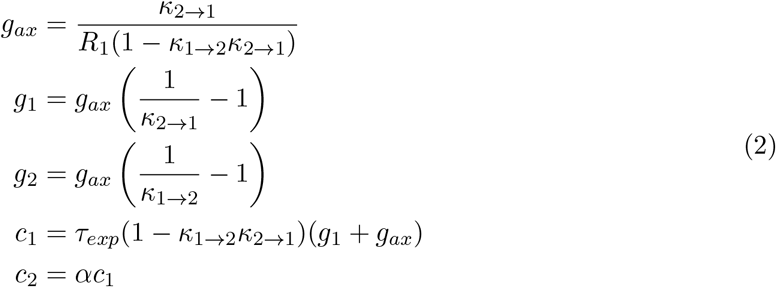

Values of these parameters throughout the coupling parameter space are shown in Fig 8A-C.

**Fig 8.**
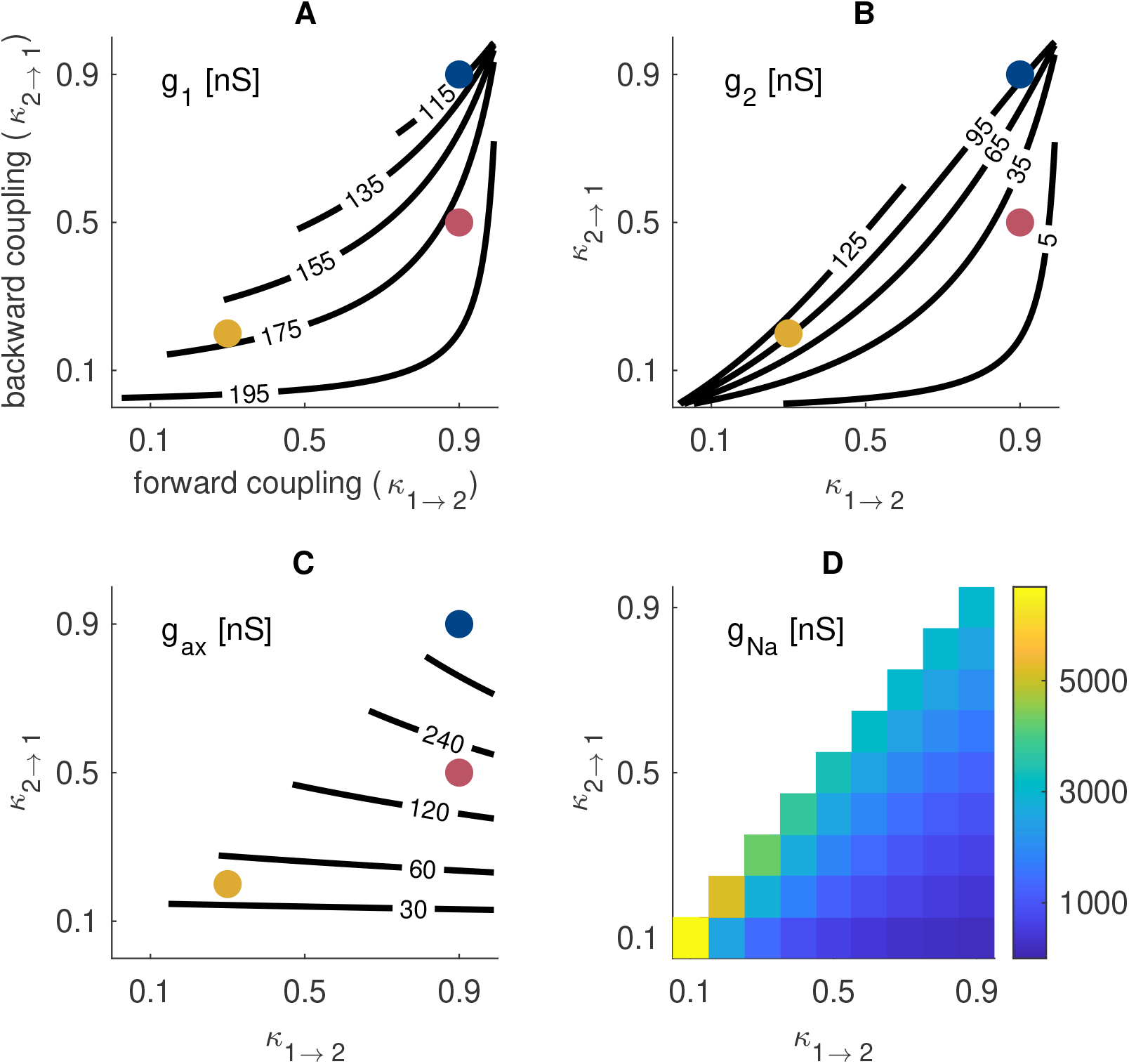
Parameter space for two-compartment NL model. (A) Total conductance in the soma compartment (*g*_1_). (B) Total conductance in the axon compartment (*g*_2_). (C) Axial conductance between the two compartments (*g*_*ax*_). Parameters in (A-C) depend uniquely on soma-axon coupling constants and commonly-reported physiological properties (input resistance and membrane time constant in the soma) and are determined for a passive model. (D) Sodium conductance in the axon compartment (*g*_*Na*_). This parameter is determined so that, at each coupling configuration, the model fired at 500 spikes/sec at ITD = 0 *µs*. Colored dots mark the specific coupling configurations used in many figures (refer to Fig 1). We used, as a reference value *κ*_1*→*2_ = 0.9 and *κ*_2*→*1_ = 0.5 since this configuration is similar to the parameter set used in [17].

### Voltage-gated spike-generating currents

The voltage-gated currents in the axon region are the spike-generating sodium current *I*_*Na*_ and the high-threshold potassium current *I*_*KHT*_. We modeled the dynamics of these currents as in [17]:

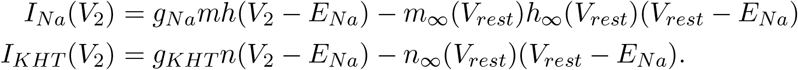

We include the second term so that these currents are zero at rest. This facilitates our exploration of coupling parameter space. This can be implemented equivalently as a shift in the leak reversal potential. Reversal potentials are *E*_*Na*_ = 35 *mV* and *E*_*K*_ = −75 *mV*.

We set maximal sodium conductance *g*_*Na*_ separately for each coupling configuration to achieve a consistent firing rate response of 500 spikes per second to in-phase synaptic inputs. We selected this criterion to be similar to the peak firing rate reported in previous modeling work [17, 30]. We found *g*_*Na*_ values to range from 200 nS to 7000 nS, roughly (Fig 8D). We set the maximal high-threshold potassium conductance to *g*_*KHT*_ = 0.3*g*_*Na*_ as in [17]. The passive leak conductance in the axon (*g*_*lk*,2_ in Eq 1) was reduced by amounts equal to *g*_*Na*_ and *g*_*KHT*_ to maintain the total axon conductance at rest determined by the initial parameter-fitting calculation (*g*_2_ in Eq 2). We did not include any voltage-gated currents in the soma (as in [18]), so leak conductance in the soma compartment is identical to *g*_1_.

The kinetics of the gating variables are governed by equations of the form

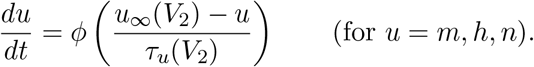

The constant *ϕ* = 4.75 adjusts for temperature at 40°C with Q10 factor 2.5. The functions *u*_*∞*_ and *τ*_*u*_ are identical to the model in [17] using the conventional definitions that *u*_*∞*_(*V*) = *α*_*u*_(*V*)*/* (*α*_*u*_(*V*) + *β*_*u*_(*V*)) and *τ*_*u*_(*V*) = 1*/* (*α*_*u*_(*V*) + *β*_*u*_(*V*)) where *α*_*u*_ and *β*_*u*_ represent opening and closing rates, respectively, for voltage-gated ion channel subunits (Table 1)

**Table 1.** Voltage-gated ion channel subunit kinetics.

|  |
| --- |
| **Na activation ($m$)** |
| $\alpha_m(V) = 3.6e^{(V+34)/7.5}$ |
| $\beta_m(V) = 3.6e^{-(V+34)/10}$ |
| **Na inactivation ($h$)** |
| $\alpha_h(V) = 0.6e^{-(V+34)/18}$ |
| $\beta_h(V) = 0.6e^{(V+34)/13.5}$ |
| **KHT activation ($n$)** |
| $\alpha_n(V) = 0.110e^{(V+19.)/9.1}$ |
| $\beta_n(V) = 0.103e^{-(V+19.)/20}$ |

### Modification of sodium inactivation for phasic model

We created a phasic version of the model by altering the steady-state function for sodium inactivation (*h*_*∞*_). The default definition of *h*_*∞*_ using the values of the *α*_*h*_ and *β*_*h*_ given in Table 1 is

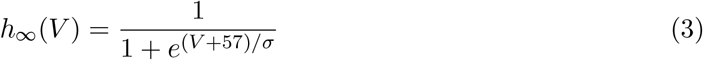

where *σ* = 7.7 reproduces the model in [17]. We found that reducing *σ* (resulting in steeper *h*_*∞*_ curve) was a practical way to toggle the model between a tonic firing mode (at the default *σ* value) and a phasic firing mode (for smaller *σ* values). We recalculated maximal sodium conductance separately for each *σ* value to maintain the consistent peak firing rate of 500 spikes/sec. It was necessary to increase *g*_*Na*_ for smaller values of *σ*. For the coupling configuration *κ*_1*→*2_ = 0.9 and *κ*_2*→*1_ = 0.5, for example, *g*_*Na*_ increased from 1240 nS for *σ* = 9 to 1838 nS for *σ* = 3.

### Synaptic current and the SAP

We modeled input currents *I*_*in*_ as either a sinusoidally-varying conductance (an idealized description of the high-frequency oscillatory input to NL neurons [17, 18, 29]) or as the summed input of simulated excitatory synaptic events. In the first case (idealized sinusoidal input), the parameter *g*_0_ is the baseline (mean) level of the input conductance and *g*_1_ is the amplitude of input oscillations. The reversal potential is *E*_*syn*_ = 0 mV for both input types. We used *f* = 4000 Hz in all simulations in this study. This value is in the range of high-frequency tones that barn owls can localize and has been used in previous modeling studies [17, 18, 30]. Our rationale for this idealized input is that tone-evoked voltage responses in the soma of NL neurons are characterized by oscillations at the tone frequency (the so-called sinusoidal analogue potential, SAP) and that SAP amplitude (not baseline level) varies with ITD [17]. The parameter *g*_1_ controls the amplitude of this idealized input with large *g*_1_ interpreted to represent preferred ITDs with coincident inputs that drive maximal firing.

For simulations in which we preferred a more biophysically-realistic description of the ITD computation performed by NL neurons, we let *I*_*in*_(*t*) = *g*(*t*)(*E*_*syn*_ - *V*_1_) where *g*(*t*) represents synaptic conductance generated by simulated trains of synaptic events filtered by short-duration excitatory post-synaptic potentials (EPSGs):

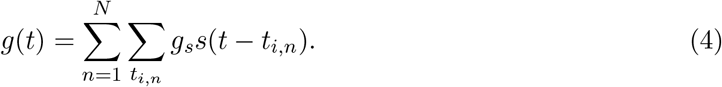

We used the synaptic model of Ashida and colleagues [29, 30] and parameters drawn from their work. Synaptic reversal potential is *E*_*syn*_ = 0 mV (as above) and the time-course of synaptic conductance *g*(*t*) is a random process constructed as the sum of unitary EPSG events produced by *N* independent input neurons whose event times *t*_*i,n*_ are sampled from an inhomogeneous Poisson process. The Poisson intensity *λ*(*t*) is the periodic function *λ*(*t*) = 2*πλ*_0_*p*_*k*_(2*πft*), where *f* = 4000 Hz is the input frequency, *λ*_0_ is the baseline rate (500 Hz), and *p*_*κ*_ is the von Mises distribution function with concentration parameter *κ* (see [29, 30] for details). The unitary EPSG events are alpha-functions

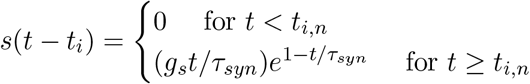

with maximal conductance *g*_0_ = 1.3 nS and time constant *τ*_*syn*_ = 40.9 *µ*s. This exceptionally brief time constant is required for the model to replicate properties of the SAP observed *in vivo* [17, 29, 30]. We used *N* = 300 for the total number of inputs (NM neurons), evenly divided into two input streams to simulate binaural (two-eared) inputs. Synaptic inputs carry ITDs when there is a time-lag between the time-courses of *λ*(*t*) used in each of the two NM population representing the two “ears.”

### Measure of coincidence detection sensitivity and ITD coding

We simulated the sound localization computation performed by NL neurons by measuring mean firing rate of the two-compartment model in response to repeated samples from the synaptic input model. Spikes (defined as an upward crossing of *V*_2_ past -30 mV were counted over a duration of 20 ms and mean spike rates were averages from 100 repetitions.

As a summary measure of ITD sensitivity we calculated the difference between in-phase and out-of-phase mean firing rates (visualized as the peak-to-trough difference in ITD tuning curve height). We denoted this statistic as Δ*R* and note that it has been used commonly used in previous studies of ITD processing including for NL neurons [17, 18, 28].

We calculated thresholds for repetitive spiking in response to sinusoidal input conductance. Repetitive firing to these inputs was defined as more than one spike in both halves of the 20 ms-long stimulus. Threshold for repetitive firing was defined as the smallest possible input strength at which repetitive firing could be observed over a range of initial values. A modified bisection search method was used so that, with *g*_0_ fixed, *g*_1_ thresholds were calculated to within ±0.5 nS. As part of the bisection search, we found it necessary to systemically test a range of initial values in order to identify the minimum threshold in cases when the model exhibited hysteresis dynamics.

### Numerical methods

Original simulation code was developed in C, python, and Matlab and is available at https://github.com/jhgoldwyn/TwoCompartmentNL. Computations to determine *g*_*Na*_ conductance values and measure ITD tuning curves were performed on a multi-CPU cluster maintained by Swarthmore College. All other computations were performed on personal laptops. Two-compartment model simulations were carried out using the forward Euler method with a 0.1 *µs* time-step. Synaptic conductance time-courses (*g*(*t*) in Eq 4) were also computed at this temporal resolution. For some coupling configurations (those with large *g*_*Na*_ values), we found it necessary to use smaller time step-sizes in the Euler calculations, in which case we linearly interpolated synaptic conductance time-courses to the smaller time steps.

### Synaptic classification by Fisher’s linear discriminant

As conceptual support for how to understand the coincidence detection computation performed by NL neurons, we considered how in-phase and out-of-phase inputs could be linearly separated (Fig 3A). Specifically, for every period of the 4 kHz stimulus, we measured the mean and amplitude (half the maximum-to-minimum range of *g*(*t*)) of synaptic input currents for one-second-long samples of the biophysically-based synaptic input model (see above). We then used Fisher’s linear discriminant to find the direction **w** along which to project these data in order to maximally separate them in-phase from out-of-phase synaptic inputs. Let **x**_*n*_ be the vector containing the mean and amplitude of *g*(*t*) on each period of an in-phase input, **y**_*n*_ the corresponding vector for out-phase inputs, and denote the means of these values (over all periods of the input) as ⟨**x**⟩ and ⟨**y**⟩, respectively. Then Fisher’s Linear Discriminant for optimally separating the in-phase and out-of-phase inputs on a cycle-by-cyle basis is to project these data onto any vector **w** in the direction of 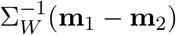 [31], where Σ_*W*_ is the within-class covariance matrix

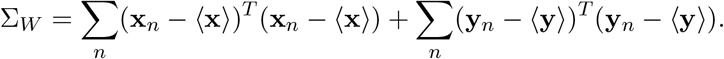

### Nonlinear integrate-and-fire model

Two features of nonlinear dynamics in the axon that we found could impact coincidence detection sensitivity are the quickness of spike initiation and the extent to which sodium current in the axon amplifies subthreshold voltages in the soma (causing the input region to deviate from linear, passive dynamics). We developed a nonlinear integrate-and-fire mode to investigate these two features. The dynamics of this model were governed by separate rules for subthreshold and suprathreshold (spike initiation) behavior to investigate these two mechanisms with direct parameter control.

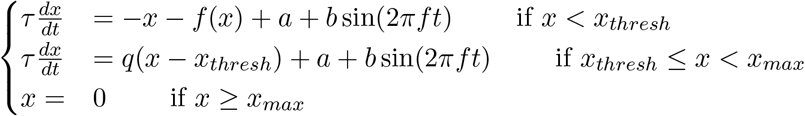

where the function *f* (*x*) has the form

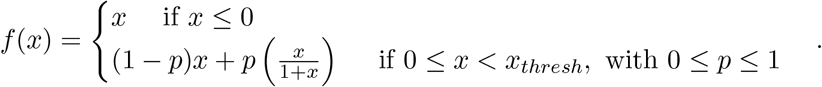

We view the piecewise-nonlinearity in *f* (*x*) as a caricature of the amplifying effect that sodium current in the axon can have if there is sufficient backpropagation from axon to soma (strong backward coupling), as shown in Fig 4A and B. In particular, the parameter *p* interpolates between linear subthreshold dynamics (*p* = 0) and supralinear subthreshold dynamics (*p* = 1).

Spike generation in the model in two phases. First, if *x*(*t*) exceeds the spike initiation threshold *x*_*thresh*_, then *x*(*t*) increases with the exponential growth rate *q*. Second, spike generation occurs at the instant at which *x*(*t*) exceeds *x*_*max*_ = 50, at which point the value of *x*(*t*) resets to *x*_*reset*_ = −5. We use the exponential growth parameter *q* to characterize the slope of spike upstroke, which we observed could change with coupling configuration (Fig 4C). The integrate-and-fire model was simulated in Matlab with code available at https://github.com/jhgoldwyn/TwoCompartmentNL.

## Acknowledgments

This work used the Strelka Computing Cluster at Swarthmore College, which is supported by the Swarthmore College Office of the Provost.

